# Three LysM effectors of *Zymoseptoria tritici* collectively disarm chitin-triggered plant immunity

**DOI:** 10.1101/2020.06.24.169789

**Authors:** Hui Tian, Craig I. MacKenzie, Luis Rodriguez-Moreno, Grardy C.M. van den Berg, Hongxin Chen, Jason J. Rudd, Jeroen R. Mesters, Bart P.H.J. Thomma

**Author notes:** Departamento de Biología Celular, Genética y Fisiología, Universidad de Málaga, Málaga, Spain (Luis Rodriguez-Moreno). These authors contributed equally (Craig I. MacKenzie, Luis Rodriguez-Moreno).

## Abstract

Chitin is a major structural component of fungal cell walls and acts as a microbe-associated molecular pattern (MAMP) that, upon recognition by a plant host, triggers the activation of immune responses. In order to avoid the activation of these responses, the Septoria tritici blotch (STB) pathogen of wheat, *Zymoseptoria tritici*, secretes LysM effector proteins. Previously, the LysM effectors Mg1LysM and Mg3LysM were shown to protect fungal hyphae against host chitinases. Furthermore, Mg3LysM, but not Mg1LysM, was shown to suppress chitin-induced reactive oxygen species (ROS) production. Whereas initially a third LysM effector gene was disregarded as a presumed pseudogene, we now provide functional data to show that also this gene encodes a LysM effector, named Mgx1LysM, that is functional during wheat colonization. While Mg3LysM confers a major contribution to *Z. tritici* virulence, Mgx1LysM and Mg1LysM contribute to *Z. tritici* virulence with smaller effects. All three LysM effectors display partial functional redundancy. We furthermore demonstrate that Mgx1LysM binds chitin, suppresses the chitin-induced ROS burst and is able to protect fungal hyphae against chitinase hydrolysis. Finally, we demonstrate that Mgx1LysM is able to undergo chitin-induced polymerisation. Collectively, our data show that *Zymoseptoria tritici* utilizes three LysM effectors to disarm chitin-triggered wheat immunity.

## INTRODUCTION

Plants deploy an effective innate immune system to recognize and appropriately respond to microbial invaders. An important part of this immune system involves the recognition of conserved microbe-associated molecular patterns (MAMPs) that are recognized by cell surface-localized pattern recognition receptors (PRRs) to activate pattern-triggered immunity (PTI) (Cook *et al.*, 2015; Jones and Dangl, 2006; Thomma *et al.*, 2001). PTI includes a broad range of immune responses, such as the production of reactive oxygen species (ROS), ion fluxes, callose deposition and defence-related gene expression (Altenbach and Robatzek, 2007; Boller and Felix, 2009; Jones and Dangl, 2006).

Chitin, a homopolymer of β-(1,4)-linked *N*-acetylglucosamine (GlcNAc), is an abundant polysaccharide in nature and a major structural component of fungal cell walls (Free, 2013). Plants secrete hydrolytic enzymes, such as chitinases, as an immune response to target fungal cell wall chitin in order to disrupt cell wall integrity, but also to release chitin molecules that act as a MAMP that can be recognized by PRRs that carry extracellular lysin motifs (LysMs) to activate further immune responses against fungal invasion (Felix *et al.*, 1993; Kombrink and Thomma, 2013; Sánchez-Vallet *et al.*, 2015). To date, chitin receptor complexes that comprise LysM-containing receptors have been characterized in Arabidopsis and rice (Cao *et al.*, 2014; Miya *et al.*, 2007; Shimizu *et al.*, 2010; Wan *et al.*, 2012). Homologs of the crucial components of these complexes have also been identified in wheat (Lee *et al.*, 2014).

In order to successfully establish an infection, fungal pathogens evolved various strategies to overcome chitin-triggered plant immunity, such as alternation of cell wall chitin in such way that it is no longer recognized (Fujikawa *et al.*, 2009; Fujikawa *et al.*, 2012), but also the secretion of effector proteins to either protect fungal cell walls against hydrolytic host enzymes or to prevent the activation of chitin-induced immunity (van den Burg *et al.*, 2006; Kombrink *et al.*, 2011; Marshall *et al.*, 2011; Mentlak *et al.*, 2012; Rovenich *et al.*, 2014; Takahara *et al.*, 2016). For example, some fungi can convert the surface-exposed chitin in fungal cell walls to chitosan, which is a poor substrate for chitinases, thus avoiding the activation of chitin-triggered immune responses during host invasion (El Gueddari *et al.*, 2002; Ride and Barber, 1990). Furthermore, from the soil-borne fungus *Verticillium dahliae* a secreted polysaccharide deacetylase was characterized to facilitate fungal virulence through direct deacetylation of chitin oligomers, converting them to chitosan that is a relatively poor inducer of immune responses (Gao *et al.*, 2019). The use of effector molecules to successfully target chitin-triggered plant immunity has been well-studied for the tomato leaf mould fungus *Cladosporium fulvum*. This fungus secretes the invertebrate chitin-binding domain (CBM14)-containing effector protein Avr4 to bind fungal cell wall chitin, resulting in the protection of its hyphae against hydrolysis by tomato chitinases (van den Burg *et al.*, 2006; van Esse *et al.*, 2007). Additionally, *C. fulvum* secretes the effector protein Ecp6 (extracellular protein 6) that carries three LysMs, binds chitin and suppresses chitin-induced plant immunity. A crystal structure of Ecp6 revealed that two of its three LysM domains undergo ligand-induced intramolecular dimerization, thus establishing a groove with ultrahigh (pM) chitin binding-affinity that enables Ecp6 to outcompete plant receptors for chitin binding (Sánchez-Vallet *et al.*, 2013). Whereas Avr4 cannot suppress chitin-triggered immunity, Ecp6 does not possess the ability to protect fungal hyphae against chitinases (Bolton *et al.*, 2008; de Jonge *et al.*, 2010). Homologs of Ecp6, coined LysM effectors, have been found in many fungi (de Jonge and Thomma, 2009). In contrast, homologs of Avr4 are less widespread (Stergiopoulos *et al.*, 2010).

*Zymoseptoria tritici* (formerly *Mycosphaerella graminicola*) is a host-specific hemibiotrophic fungus and the causal agent of Septoria tritici blotch (STB) of wheat (*Triticum* spp.) (Eyal, 1999). Upon infection, wheat plants undergo an extended period of symptomless colonization of approximately one week, followed by the death of host tissues coinciding with rapid invasive growth and asexual reproduction of the fungus (Glazebrook, 2005; Kema *et al.*, 1996; Pnini-Cohen *et al.*, 2000). This transition from biotrophic to necrotrophic growth of *Z. tritici* is associated with the induction of host immune processes such as a hypersensitive response (HR)-like programmed cell death and differential expression of wheat mitogen-activated protein kinase (MAPK) genes (Rudd *et al.*, 2008). Three LysM effector genes were previously identified in the *Z. tritici* genome (Marshall *et al.*, 2011). These comprise *Mg1LysM* and *MgxLysM* that encode LysM effector proteins that carry a single LysM only, and *Mg3LysM* encoding an effector with three LysMs (Marshall *et al.*, 2011). Whereas *Mg1LysM* and *Mg3LysM* were subjected to functional analysis, *MgxLysM* was disregarded because this gene lacked expressed sequence tag (EST) support and was believed to contain an intronic repeat insertion, rendering it a pseudogene. Both *Mg1LysM* and *Mg3LysM* were found to be induced during wheat infection, and both proteins were found to bind chitin. However, only Mg3LysM was found to suppress chitin-induced plant immunity (Marshall *et al.*, 2011). Surprisingly, and in contrast to Ecp6, both Mg1LysM and Mg3LysM were found to protect fungal hyphae against plant chitinase activity. Recently, a crystal structure was generated and revealed that Mg1LysM undergoes chitin-dependent dimerization of ligand-independent homodimers, and it was proposed that chitin-induced polymerization of Mg1LysM in the fungal cell wall confers protection against chitinases (Sánchez-Vallet *et al.*, 2020). However, thus far the mechanism underlying the protection of cell walls by Mg3LysM remains unclear. In this study, we revisit the previously discarded *MgxLysM* gene and evaluate its contribution to *Z. tritici* virulence on wheat plants.

## RESULTS

### *Mgx1LysM* is expressed during wheat colonization

Although *MgxLysM* was previously reported to be a pseudogene and found not to be induced upon wheat infection (Marshall *et al.*, 2011), a more recent transcriptome profiling study on wheat demonstrated *MgxLysM* expression during host colonization, demonstrating that the initial assessment was incorrect (Rudd *et al.*, 2015). Thus, we propose to rename MgxLysM to Mgx1LysM, according to the single LysM domain in the protein, similar to the previously described Mg1LysM effector (Marshall *et al.*, 2011).

To confirm the expression of *Mgx1LysM* in *Z. tritici* upon host colonization, we inoculated the wild-type strain IPO323 onto wheat leaves and sampled leaves at 0, 4, 8, 10 and 14 days post inoculation (dpi). In addition, we subjected IPO323 growing *in vitro* in Czapek-Dox broth (CDB) and in potato dextrose broth (PDB) to expression analysis. We confirmed that *Mgx1LysM* is not expressed upon growth *in vitro*, but only during host colonization at all tested time points (Fig. 1). More specifically, *Mgx1LysM* expression was strongly induced at 4 dpi, peaked at 8 dpi and dramatically decreased by 10 dpi. Interestingly, the peak of expression at 8 dpi is around the transition time when the infection switches from asymptomatic to symptomatic with the appearance of lesions on wheat leaves (Marshall *et al.*, 2011).

**Fig. 1.**
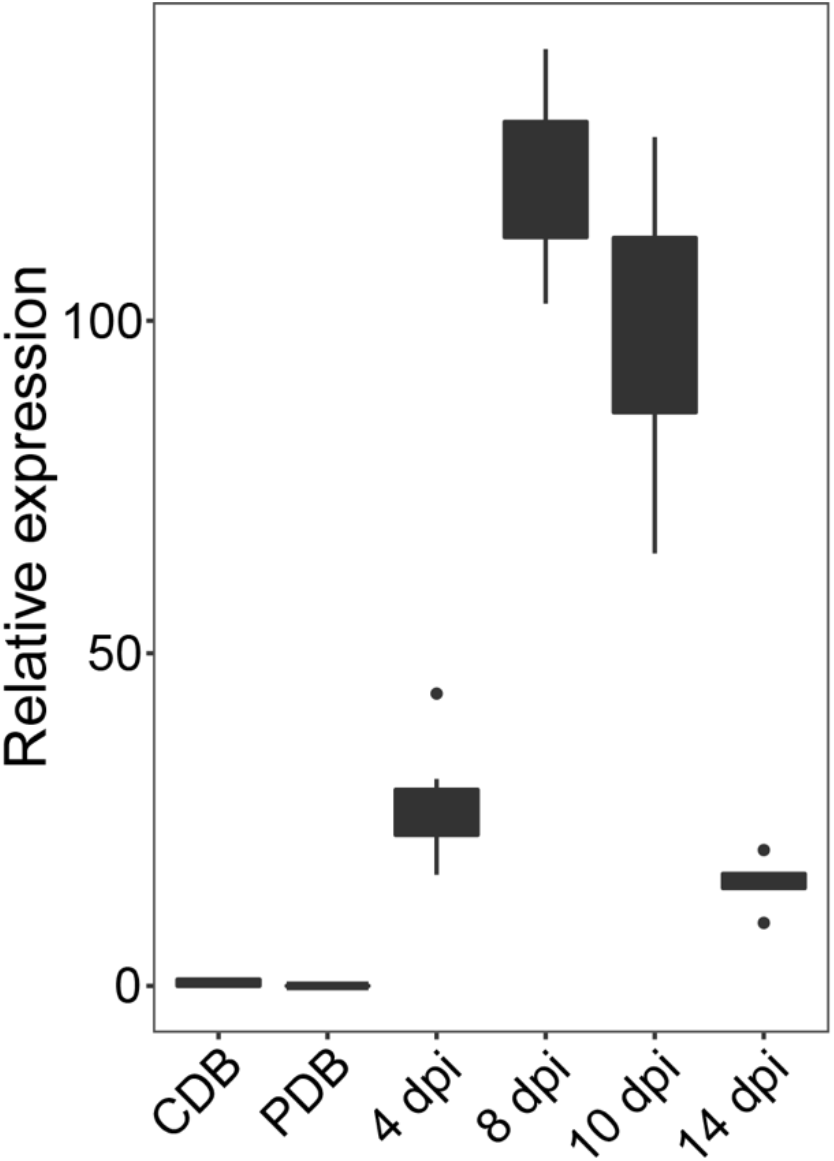
Expression of *Mgx1LysM* is induced in *Zymoseptoria tritici* upon inoculation on wheat plants. Relative expression of *Mgx1LysM* at 4, 8, 10 and 14 days post inoculation on wheat plants and upon growth *in vitro* in Czapek-dox (CDB) or potato dextrose broth (PDB) when normalized to *Z. tritici β-tublin*. The boxplot was made with RStudio using the package ggplot2.

### Mgx1LysM contributes to *Z. tritici* virulence on wheat and displays functional redundancy with Mg1LysM and Mg3LysM

Since *Mgx1LysM* is expressed by *Z. tritici* during colonization of wheat plants, we further assessed whether Mgx1LysM contributes to *Z. tritici* virulence and whether it shares functional redundancy with Mg1LysM and Mg3LysM. To this end, we inoculated the single-gene deletion mutants *ΔMg1, ΔMgx1* and *ΔMg3*, the double-gene deletion mutants *ΔMg1-ΔMgx1*, *ΔMgx1*-*ΔMg3* and *ΔMg1-ΔMg3*, and the triple-gene deletion mutant *ΔMg1-ΔMgx1*-*ΔMg3*, all of which were generated in a *Δku70* mutant, onto wheat plants. The *Δku70* mutant is with unaltered virulence but improved homologous recombination frequencies (Bowler *et al.*, 2010), and was used as wild-type (WT) in this study. By 17 days post inoculation (dpi), the WT strain caused typical necrosis symptoms on the wheat leaves, while the *ΔMg3* strain caused much less necrotic symptoms (Fig. 2AB) as previously reported (Marshall *et al.*, 2011). Furthermore, like previously reported, plants inoculated with the *ΔMg1* strain developed similar necrosis as with the WT strain. We now show not only that the *ΔMgx1* strain caused similar levels of necrosis as the WT and *ΔMg1* strain, but also that the *ΔMg1*-*ΔMgx1* strain shows no apparent decrease in disease development, suggesting that these two LysM effectors are dispensable for virulence of *Z. tritici*. In line with these observations, both the *ΔMgx1*-*ΔMg3* and *ΔMg1*-*ΔMg3* strains induced similar symptoms as the *ΔMg3* strain (Fig. 2AB). Nevertheless, the necrotic symptoms caused by inoculation with the *ΔMg1*-*ΔMgx1*-*ΔMg3* strain were drastically reduced when compared with those caused by the *ΔMg3* strain. Collectively, these findings suggest that Mg3LysM is the most important LysM effector for *Z. tritici* disease development, and that Mgx1LysM and Mg1LysM contribute to disease development through redundant functionality.

**Fig. 2.**
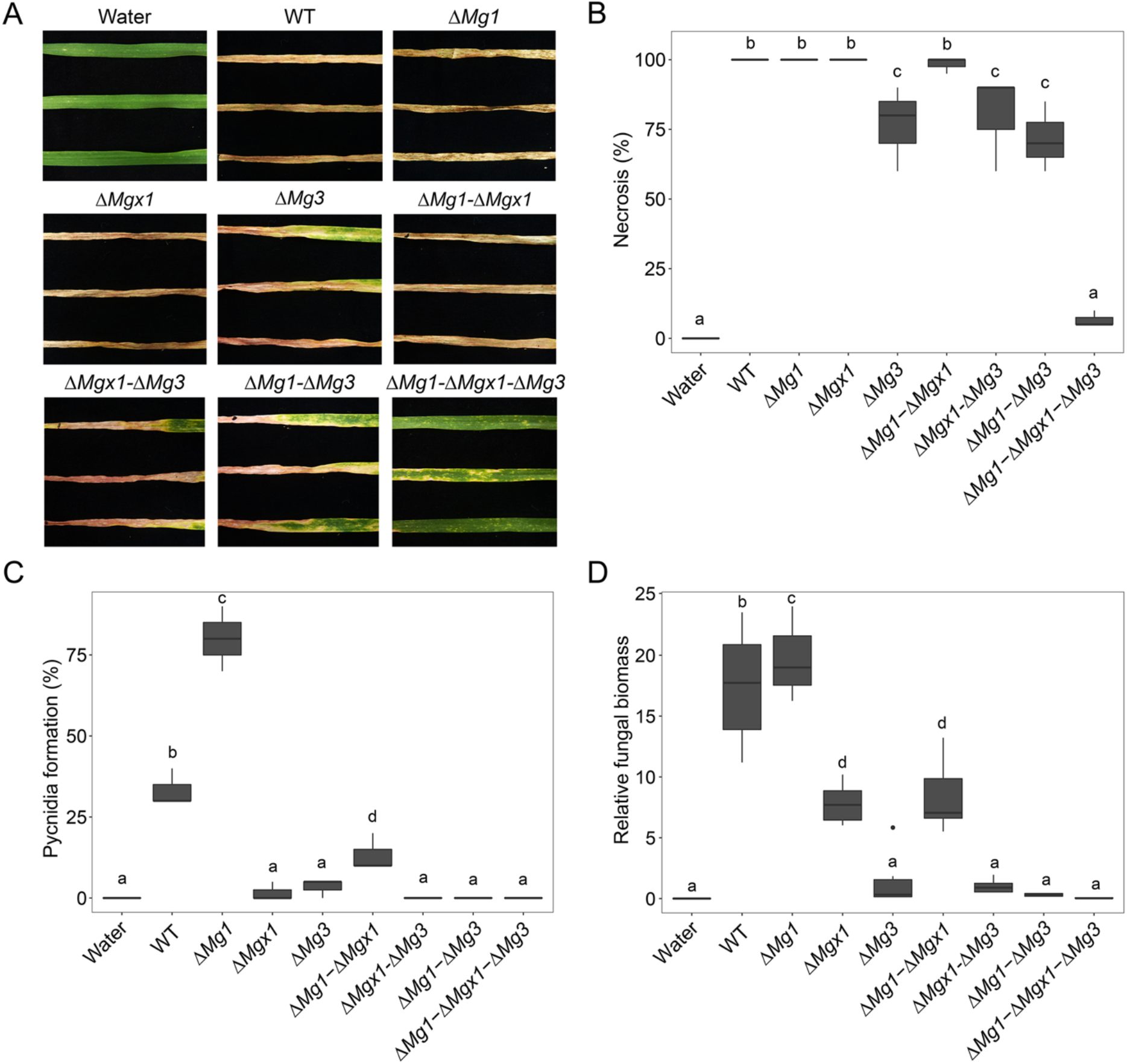
Mgx1LysM contributes to *Z. tritici* virulence on wheat and displays functional redundancy with Mg1LysM and Mg3LysM. (A) Disease symptoms on wheat leaves at 17 days post inoculation (dpi) with the wild-type strain (WT) and LysM effector gene deletion strains. Quantification of the necrotic area (B) and of the area displaying the formation of asexual fruiting bodies (pycnidia) (C) on wheat leaves inoculated with WT and LysM effector gene deletion strains at 17 dpi. (D) Fungal biomass determined with real-time PCR on *Z. tritici β-tubulin* relative to the wheat cell division control gene, on wheat leaf samples harvested at 17 dpi. Graphs were made with RStudio using the package of ggplot2 and different letters indicating significant differences between each inoculation were calculated with IBM Statistics 25 with One-way ANOVA (Duncan; P<0.05). Fungal inoculation experiments were conducted on six plants with six first-primary leaves per inoculation and repeated three times with similar results.

To further substantiate our findings, we also determined the formation of asexual fruiting bodies (pycnidia) as a measure for fungal colonization on the wheat leaves at 17 dpi. To this end, we determined the percentage of leaf surface displaying pycnidia. Surprisingly, repeated assays revealed that the *ΔMg1* strain developed significantly more pycnidia than the WT strain, whereas the *ΔMgx1* strain, like the *ΔMg3*, produced no to only a few pycnidia (Fig. 2C). Accordingly, whereas the *ΔMg1*-*ΔMgx1* strain developed an intermediate number of pycnidia, all mutants that involved *ΔMg3* were devoid of pycnidia (Fig. 2C). These data first of all suggest that symptom development does not correlate with fungal colonization levels as measured by pycnidia formation and, furthermore, that the three LysM effectors display differential roles in fungal colonization.

To further substantiate the fungal colonization assessments, we measured fungal biomass with real-time PCR. While the *ΔMg1* strain developed a similar amount of fungal biomass as the WT strain, both the *ΔMgx1* and *ΔMg1*-*ΔMgx1* strains displayed significantly compromised colonization, but not as compromised as the *ΔMg3* strain or the double mutants and triple mutant that carry *ΔMg3* (Fig. 2D). These observations confirm the discrepancy between fungal colonization and symptom development, and also fit with the differential contribution of the LysM effectors to fungal colonization. Consequently, our data reveal that the three LysM effectors make differential contributions to symptom display, that is accompanied by distinct differential contributions to fungal colonization. Thus, our findings present evidence for partially redundant, but also partially divergent, contributions of the three LysM effectors to *Z. tritici* virulence.

### Mgx1LysM binds chitin and suppresses the chitin-induced ROS burst

To investigate how Mgx1LysM contributes to *Z. tritici* virulence during wheat colonization, we first assessed its substrate-binding characteristics. Mgx1LysM was heterologously expressed in *E. coli* and subjected to a polysaccharide precipitation assay. Mgx1LysM was incubated with chitin beads and shrimp shell chitin, but also with plant-derived cellulose and xylan, revealing that Mgx1LysM binds chitin beads and shrimp shell chitin but not cellulose or xylan (Fig. 3). Thus, Mgx1LysM resembles Mg1LysM that similarly binds chitin but not cellulose or xylan (Fig. 3).

**Fig. 3.**
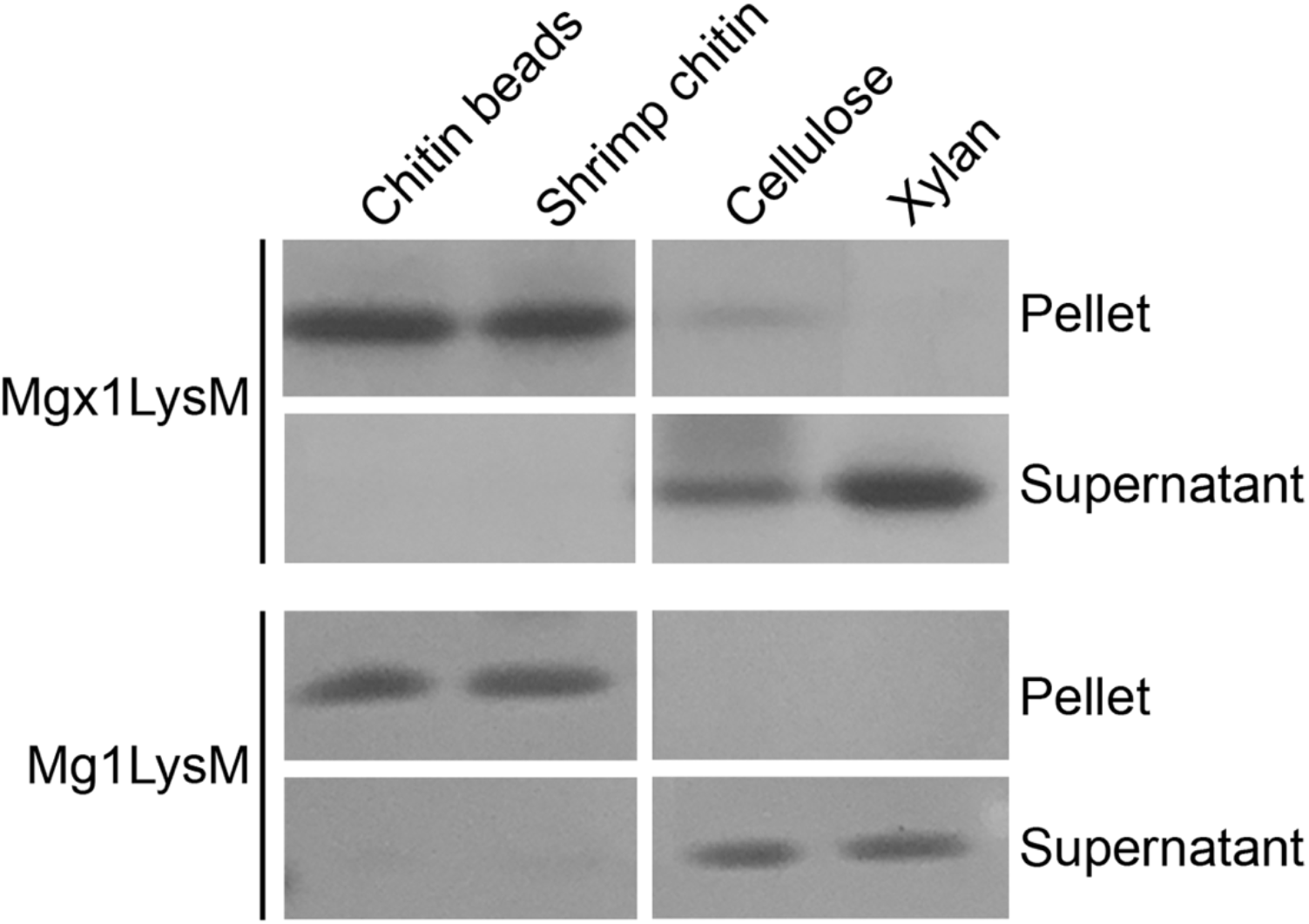
Mgx1LysM binds chitin. *E. coli*-produced Mgx1LysM and Mg1LysM were incubated with four chitin products for 6 hours and, after centrifugation, pellets and supernatants were analysed using polyacrylamide gel electrophoresis followed by CBB staining.

To test whether Mgx1LysM can prevent chitin-triggered immunity in plants, the occurrence of a chitin-induced ROS burst was assessed in *Nicotiana benthamiana* leaf discs upon treatment with 10 μM chitohexaose (chitin) in the presence or absence of effector protein. As previously demonstrated (de Jonge *et al.*, 2010), *C. fulvum* Ecp6 suppresses ROS production in this assay (Fig. 4). Remarkably, pre-incubation of 10 μM chitin with 50 μM Mgx1LysM prior to the addition to leaf discs led to a significant reduction of the ROS burst (Fig. 4), demonstrating its ability to suppress chitin-induced plant immune responses. This finding was unexpected because we previously found that its close homolog Mg1LysM cannot suppress a chitin-induced defense response in a tomato cell culture (Marshall *et al.*, 2011), albeit that in that study Mg1LysM was heterologously produced in the yeast *Pichia pastoris* rather than in *E. coli*. To revisit this initial observation, we now test whether *E. coli*-produced Mg1LysM is able to suppress the chitin-induced ROS burst. Indeed, similar to the results obtained for Mgx1LysM, we observed that pre-incubation of 10 μM chitin with 50 μM Mg1LysM prior to the addition to leaf discs led to a significantly compromised ROS burst. Thus, both LysM effectors can suppress chitin-triggered host immunity.

**Fig. 4.**
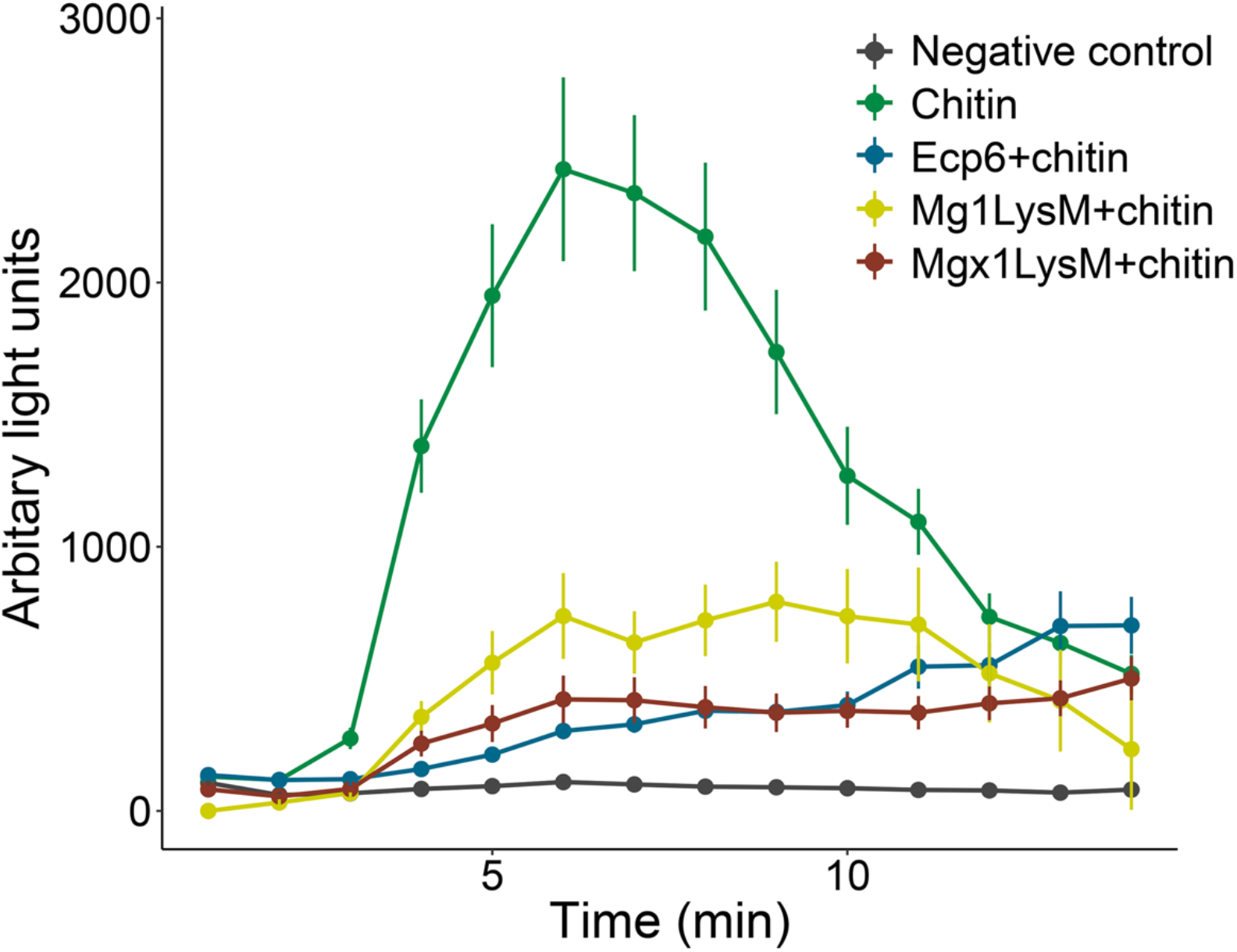
Mgx1LysM suppresses the chitin-induced reactive oxygen species (ROS) burst. Leaf discs of *Nicotiana benthamiana* were treated with chitohexaose (chitin) to induce ROS production. Chitin was pre-incubated with Ecp6, Mg1LysM or Mgx1LysM for two hours and subsequently added to the leaf discs. Error bars represent standard errors from five biological replicates.

### Mgx1LysM protects hyphae against chitinases

We previously demonstrated that Mg1LysM can protect fungal hyphae against chitinase hydrolysis (Marshall *et al.*, 2011). To evaluate a possible role in hyphal protection, Mgx1LysM was tested for its ability to protect hyphae of *Trichoderma viride*, a fungus that exposes its cell wall chitin *in vitro*, against chitinases was tested (Mauch *et al.*, 1988). The *C. fulvum* effector protein Avr4 and Mg1LysM were used as positive controls based on their previously demonstrated ability to protect fungal hyphae (van den Burg *et al.*, 2006; Marshall *et al.*, 2011). As expected, while the addition of chitinase drastically inhibited *T. viride* hyphal growth, Avr4 as well as Mg1LysM protected the hyphae against chitinase hydrolysis (Fig. 5). Furthermore, Mgx1LysM similarly protected the hyphae against chitinase hydrolysis (Fig. 5).

**Fig. 5.**
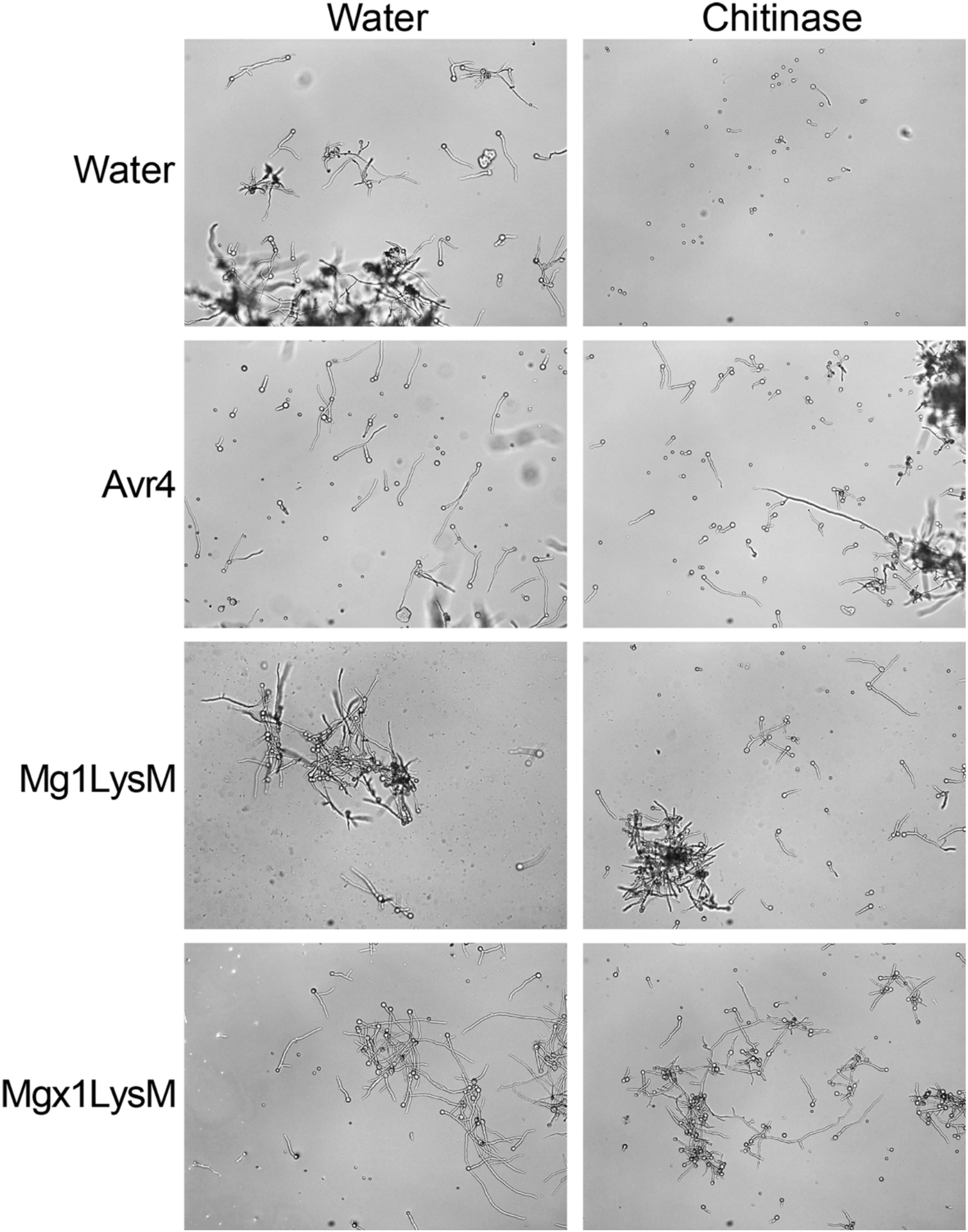
Mgx1LysM protects hyphal growth of *Trichoderma viride* against chitinase hydrolysis. Microscopic pictures of *T. viride* grown *in vitro* with or without two hours of preincubation with *C. fulvum* Avr4, or *Z. tritici* Mg1LysM or Mgx1LysM, followed by the addition of chitinase or water. Pictures were taken ~4 hours after the addition of chitinase.

### Mgx1LysM undergoes chitin-dependent polymerization

Recently, Mg1LysM was demonstrated to protect fungal hyphae through chitin-dependent polymerization of chitin-independent Mg1LysM homodimers (Sánchez-Vallet *et al.*, 2020). To assess whether this trait is shared by Mgx1LysM, the amino acid sequence of Mgx1LysM was aligned with Mg1LysM, displaying an overall sequence identity of 44% (Fig. 6A). As expected, the predicted three-dimensional structure of Mgx1LysM shows a typical LysM fold with two antiparallel β-sheets adjacent to two α-helices (Fig. 6B) (Bateman and Bycroft, 2000; Bielnicki *et al.*, 2006; Liu *et al.*, 2012; Sánchez-Vallet *et al.*, 2013; Sánchez-Vallet *et al.*, 2020). More importantly, similar to Mg1LysM, Mgx1LysM carries a relatively long N-terminal sequence (Fig. 6B). For Mg1LysM it was recently shown that this N-terminal tail of a single monomer runs antiparallel with the tail of another Mg1LysM monomer, leading to the formation of ligand-independent homodimers (Sánchez-Vallet *et al.*, 2020). Structural modelling of Mgx1LysM suggests that this LysM effector is also able to dimerize via its N-terminal tail (Fig. 6B).

**Fig. 6.**
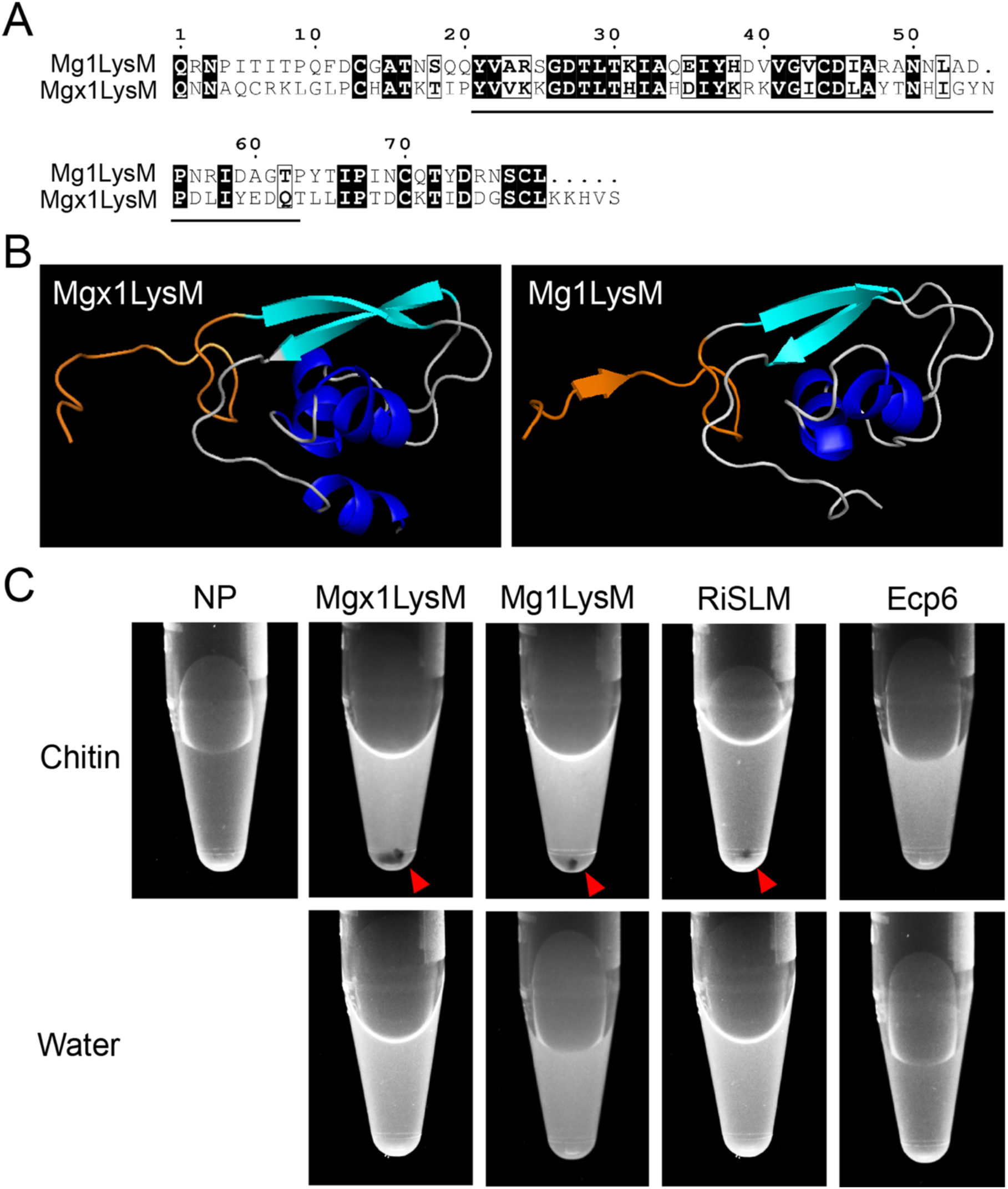
Mgx1LysM undergoes chitin-induced polymerization. (A) Amino acid sequence alignment of Mgx1LysM and Mg1LysM. The LysM is indicated with black underlining. (B) I-TASSER software-based *in-silico* prediction of the three-dimensional structure of Mgx1LysM (left) based on the recently generated crystal structure of Mg1LysM (right) (Sánchez-Vallet et al., 2020). The *N*-terminal 15 amino acids of both proteins are depicted in orange. Structures are visualized using the PyMOL molecular graphics system (Schrodinger LLC, 2015). (C) The LysM effector Mgx1LysM, together with RiSLM and Mg1LysM as positive controls, and Ecp6 as negative control, were incubated with chitohexaose (chitin) or water. After overnight incubation, methylene blue was added and protein solutions were centrifuged, resulting in protein pellets (red arrowheads) as a consequence of polymerization for Mgx1LysM, Mg1LysM and RiSLM, but not for Ecp6.

Besides ligand-independent dimerization, the crystal structure of Mg1LysM furthermore revealed chitin-dependent dimerization of the ligand-independent homodimers (Sánchez-Vallet *et al.*, 2020). Based on further biochemical evidence the occurrence of chitin-induced polymeric complexes was demonstrated (Sánchez-Vallet *et al.*, 2020). Thus, to assess whether Mgx1LysM similarly undergoes chitin-induced polymerization, a centrifugation assay was performed using *E. coli*-produced Mgx1LysM, while Mg1LysM and RiSLM were included as positive controls and Ecp6 was used as negative control. After incubation with chitin, protein samples were centrifuged at 20,000 g in the presence of 0.002% methylene blue to visualize the protein. Similar as for Mg1LysM and RiSLM, a clear Mgx1LysM pellet emerged upon incubation with chitin, and not in the absence of chitin, while no chitin-induced polymerisation was observed for Ecp6 (Fig. 6C). Collectively, our data indicate that Mgx1LysM, like Mg1LysM, undergoes not only chitin-dependent dimerization, but also ligand-independent dimerization through interactions at the N-termini of Mgx1LysM monomers, leading to polymerization of the LysM effector protein in the presence of chitin.

## DISCUSSION

In this study, we demonstrate that the previously disregarded LysM effector gene as a presumed pseudogene of the fungal wheat pathogen *Z. tritici*, *Mgx1LysM*, is a functional LysM effector gene that plays a role in *Z. tritici* virulence during infection of wheat plants. Like the previously characterized *Z. tritici* LysM effectors Mg1LysM and Mg3LysM, Mgx1LysM binds chitin (Fig. 3), suppresses chitin-induced ROS production (Fig. 4) and can protect fungal hyphae against chitinase hydrolysis (Fig. 5). Moreover, like Mg1LysM, Mgx1LysM polymerizes in the presence of chitin (Fig. 6). Through these activities, Mgx1LysM makes a noticeable contribution to *Z. tritici* virulence on wheat plants (Fig. 2).

Merely based on expression profile as well as biological activities, the three genes seem to behave in an similar fashion and complete redundancy could be expected. However, this is not what we observed in the mutant analyses, as these revealed that Mg3LysM confers the largest contribution, as targeted deletion of *Mg3LysM*, but not of Mg1LysM or Mgx1LysM, results in a noticeable difference in symptomatology. Moreover, even the simultaneous deletion of *Mgx1LysM* and *Mg1LysM* did not lead to compromised necrosis development, although deletion of these two genes from the *Mg3LysM* deletion strain in the triple mutant resulted in a further decrease of virulence. Thus, although it can be concluded that all three LysM effectors contribute to fungal virulence, these findings are suggestive of partially redundant and partially additive activities. This suggestion is further reinforced when assessing pycnidia development and fungal colonization data that demonstrate that single LysM effector deletions have significant effects on these traits. However, it presently remains unknown through which functional divergence these differential phenotypes are established.

The ability to protect fungal hyphae against chitinase hydrolysis that is shared by the three *Z. tritici* LysM effectors (Fig. 5) has previously been recorded for some, but not all, LysM effectors from other fungal species as well. For example, although *Verticillium dahliae* Vd2LysM and *Rhizophagus irregularis* RiSLM can protect hyphae as well (Kombrink *et al.*, 2017; Zeng *et al.*, 2020), *C. fulvum* Ecp6, *Colletotrichum higginsianum* ChElp1 and ChElp2, and *Magnaporthe oryzae* MoSlp1 do not possess such activity (de Jonge *et al.*, 2010; Mentlak *et al.*, 2012; Takahara *et al.*, 2016). Intriguingly, all LysM effectors that contain a single LysM characterized to date (Mg1LysM, Mgx1LysM, RiSLM) were found to protect fungal hyphae. However among the ones with two LysM domains members are found that do (Vd2LysM) and that do not (ChElp1, ChElp2, MoSlp1) protect, which is also true for members with three LysMs (Mg3LysM versus Ecp6, respectively), suggesting that the ability to protect hyphae is not determined by the number of LysMs in the effector protein. Previously, a mechanistic explanation for the ability to protect fungal cell wall chitin has been provided for the *C. fulvum* effector protein Avr4 that acts as a functional homolog of LysM effectors that protect fungal hyphae, but that binds chitin through an invertebrate chitin-binding domain (CBM14) rather than through LysMs (van den Burg *et al.*, 2006). Intriguingly, Avr4 strictly interacts with chitotriose, but binding of additional Avr4 molecules to chitin occurs through cooperative interactions between Avr4 monomers, which can explain the effective shielding of cell wall chitin (van den Burg *et al.*, 2004). Despite being a close relative of *C. fulvum* in the Dothidiomycete class of Ascomycete fungi, *Z. tritici* lacks an Avr4 homolog (Stergiopoulos *et al.*, 2010). This may explain why the *Z. tritici* LysM effectors, in contrast to *C. fulvum* Ecp6, evolved the ability to protect fungal cell wall chitin. Recently, it has been proposed that the hyphal protection by LysM effectors that contain only a single LysM, including Mg1LysM and RiSLM, is due to chitin-induced polymerisation, leading to contiguous LysM effector filaments that are anchored to chitin in the fungal cell wall to protect these cell walls (Sánchez-Vallet *et al.*, 2020). Here, we show that Mgx1LysM similarly undergoes chitin-induced polymer formation (Fig. 6).

It was previously reported that Mg1LysM was incapable of suppressing chitin-induced immune responses (Marshall *et al.*, 2011), in contrast to the immune-suppressive activity of Mg3LysM. A mechanistic explanation for this observation was found in the observation that Ecp6, being a close homolog of Mg3LysM, was able to efficiently sequester chitin oligomers from host receptors through intramolecular LysM dimerization, leading to a binding groove with ultra-high chitin-binding affinity. As a single LysM-containing effector protein, Mg1LysM lacks the ability to undergo intramolecular LysM dimerization, and thus to form an ultra-high affinity groove for chitin binding, which could explain the inability to suppress immune responses by out-competition of host receptor molecules for chitin binding. However, this mechanistic explanation was recently challenged by data showing that the *R. irregulars* RiSLM is able to suppress chitin-triggered immunity as well (Zeng *et al.*, 2020). In the present study we show that not only Mgx1LysM can suppress chitin-triggered immunity, but also that Mg1LysM possesses this activity (Fig. 4). However, it needs to be acknowledged that whereas we used *P*. *pastoris*-produced protein in our initial analyses (Marshall *et al.*, 2011), we used *E. coli*-produced protein in the current study. More recent insights after the publication of our initial study have revealed that LysM effector proteins may bind chitin fragments that are released from the *P. pastoris* cell walls during protein production, which may compromise the activity of the protein preparation in subsequent assays (Kombrink *et al.*, 2017; Sánchez-Vallet *et al.*, 2013; Sánchez-Vallet *et al.*, 2020). As the *E. coli* cell wall is devoid of chitin, partially or fully inactive protein preparations due to occupation of the substrate binding site are unlikely to occur. However, since Mg1LysM, Mgx1LysM and RiSLM are able to suppress chitin-triggered immunity, a mechanistic explanation needs to be provided for the suppressive activity that does not involve substrate sequestration purely based on chitin-binding affinity. Possibly, these LysM effectors are able to perturb the formation of active chitin receptor complexes by binding to receptor monomers in a similar fashion as has been proposed for LysM2 of Ecp6 (Sánchez-Vallet *et al.*, 2013; Sánchez-Vallet *et al.*, 2015) to prevent the activation of chitin-triggered immune responses. Alternatively, precipitation of polymeric complexes formed by LysM effectors and released chitin oligosaccharides may provide a mechanism to eliminate these oligosaccharides and prevent their interaction with host receptor molecules.

## MATERIALS AND METHODS

### Gene expression analysis

Total RNA was isolated using the RNeasy Plant Mini Kit (Qiagen, Maryland, USA). For each sample, 2 μg RNA was used for cDNA synthesis with M-MLV Reverse Transcriptase (Promega, Madison, USA) and 1 μL of the obtained cDNA was used for real-time PCR with SYBR™ green master mix (Bioline, Luckenwalde, Germany) on a C1000 Touch™ Thermal Cycler (Bio-Rad, California, USA). Expression of *Mgx1LysM* was normalized to the *Z. tritici* housekeeping gene *β-tubulin* using primer pairs *Mgx1LysM*-F/*Mgx1LysM*-R and *Ztβtubulin*-F/R, respectively (Table S1). Relative expression was calculated with the E^−ΔCt^ method and the boxplot was made with RStudio using the package of ggplot2 (R Core Team, 2017; Wickham, 2016).

### Heterologous protein production in *E. coli*

Signal peptide prediction was performed using SignalP 5.0 (http://www.cbs.dtu.dk/services/SignalP/). The coding region for the mature Mgx1LysM protein was amplified from *Z. tritici* IPO323 genomic cDNA using primers *Mgx1LysM*-cDNA-F/ R (Table S1) and cloned into the pETSUMO vector and transformed as pETSUMO-Mgx1LysM into *E. coli* strain Origami for heterologous protein production as a fusion protein with a 6×His-SUMO affinity-tag. *Mgx1LysM* expression was induced with 0.2 mM isopropyl β-D-1-thiogalactopyranoside (IPTG) at 28°C overnight. Next, *E. coli* cells were harvested by centrifugation at 3,800 g for one hour and resuspended in 20 mL cell lysis buffer (50 mM Tris-HCl pH 8.5, 150 mM NaCl, 2 mL glycerol, 120 mg lysozyme, 40 mg deoxycholic acid, 1.25 mg DNase I and 1 protease inhibitor pill) and incubated at 4°C for two hours with stirring, and centrifuged at 20,000 g for one hour. The resulting cleared supernatant was immediately placed on ice and subjected to further purification.

The His60 Ni Superflow Resin (Clontech, California, USA) was used for Mgx1LysM purification and first equilibrated with wash buffer (50 mM Na_2_HPO_4_, 150 mM NaCl, 10 mM imidazole, pH 8.0) after which the protein preparation was loaded on the column. The target protein was eluted with elution buffer (50 mM Na_2_HPO_4_, 150 mM NaCl, 300 mM imidazole, pH 8.0), and purity of the elution was tested on an SDS-PAGE gel followed by Coomassie brilliant blue (CBB) staining. The 6×His-SUMO affinity-tag was cleaved with the SUMO Protease ULP1 during overnight dialysis against 200 mM NaCl. Non-cleaved Mgx1LysM fusion protein was removed using His60 Ni Superflow resin, and the flow-through with cleaved Mgx1LysM was adjusted to the required concentration.

### Chitin binding assay

*E. coli*-produced proteins were adjusted to a concentration of 30 μg/mL in chitin binding buffer (50 mM Tris PH 8.0, 150 mM NaCl) and 800 μL of protein solution was incubated with 50 μL of magnetic chitin beads, or 5 mg of crab shell chitin, cellulose or xylan in a rotary shaker at 4°C for 6h. The insoluble fraction was pelleted by centrifuging at 13,500 g for 5 min and resuspend in 100 μL demineralized water. Supernatants were collected into Microcon Ultracel YM-10 tubes (Merck, Darmstadt, Germany) and concentrated a volume of approximately 100 μL. For each of the insoluble carbohydrates, 30 μL of the pellet solution and the concentrated supernatant was incubated with 10 μL of SDS-PAGE protein loading buffer (4×; 200 mM Tris-HCl, pH 6.5, 0.4 M dithiothreitol, 8 % sodium dodecyl sulfate, 6 mM bromophenol blue, 40 % glycerol) and incubated at 95°C for 10 min. Samples were loaded into an SDS-PAGE gel followed by CBB staining.

### Hyphal protection against chitinase hydrolysis

*Trichoderma viride* conidiospores were harvested from five-day-old potato dextrose agar (PDA; OXOID, Basingstoke, United Kingdom), washed with sterile water, and adjusted to a concentration of 10^6^ spores/mL with potato dextrose broth (PDB; Becton Dickinson, Maryland, USA). Conidiospore suspensions were dispensed into a 96-well microtiter plate in aliquots of 50 μL and incubated at room temperature overnight. Effector proteins were added to a final concentration of 10 μM, and after 2 h of incubation 3 μL of chitinase from *Clostridium thermocellum* (Creative Enzymes, New York, USA) was added into the appropriate wells. As control, sterile water was added. All treatments were further incubated for 4 h and hyphal growth was inspected with a Nikon H600L microscope.

### Reactive oxygen species measurement

Reactive oxygen species (ROS) production measurements were performed using three *Nicotiana benthamiana* leaf discs (Ø = 0.5 cm) per treatment, which were collected from two-week-old *N. benthamiana* plants, placed into a 96-well microtiter plate, and rinsed with 200 μL demineralized water. After 24 hours the water was replaced by 50 μL fresh demineralized water and the plate was incubated for another hours at room temperature. Meanwhile, mixtures of (GlcNAc)_6_ (IsoSep AB, Tullinge, Sweden) and effector proteins were incubated for two hours. In total, 20 μL of (GlcNAc)_6_ was added in a final concentration of 10 μM to trigger ROS production in the absence or presence of 100 μL of effector protein in a final concentration of 50 μM in measuring solution containing 100 μM L-012 substrate (FUJIFILM, Neuss, Germany) and 40 μg/mL horseradish peroxidase (Sigma-Aldrich, Missouri, USA). Chemiluminescence measurements were taken every minute over 30 min in a CLARIOstar microplate reader (BMG LABTECH, Ortenberg, Germany).

### *Agrobacterium tumefaciens*-mediated *Z. tritici* transformation

To generate *Mgx1LysM* deletion mutants, approximately 1.0 kb upstream and 1.2 kb downstream fragments of *Mgx1LysM* were amplified from genomic DNA of *Z. tritici* IPO323 using primer pairs *Mgx1LysM*-userL-F/R and *Mgx1LysM*-userR-F/R (Table S1) and the amplicons were cloned into vector pRF-NU2 as previously described (Frandsen et al, 2008). The resulting deletion construct was transformed into *Z. tritici* mutant *Δku70* and the previously generated *ΔMg1LysM*, *ΔMg3LysM* and *ΔMg1*-*ΔMg3* to generate double- and triple-gene deletion mutants. In short, minimal medium (MM) and induction medium (IM) were prepared at a pH of 7.0 and *Z. tritici* conidiospores were collected, washed and adjusted to a final concentration of 10^7^ spores/mL. Transformation plates were incubated at 16°C in dark for two to three weeks. Putative transformants were transferred to PDA plates supplemented with 200 μg/mL cefotaxime and 25 μg/mL nourseothricin (Sigma-Aldrich, Missouri, USA) and absence of *Mgx1LysM* was confirmed with PCR using the gene-specific primers *Mgx1LysM*-F/*Mgx1LysM*-R and the primer pair with NAT-F as the forward primer that targets the nourseothricin cassette and the reverse primer *Mgx1LysM*-out-R targeting the downstream fragment of *Mgx1LysM* (Table S1).

### *Z. tritici* inoculations on wheat

For all inoculation assays, the wheat cultivar “Riband” was used. *Z. tritici* wild-type strain IPO323 and the mutants were grown either on yeast extract peptone dextrose (YPD; 10 g yeast extract/L, 20 g peptone/L, 20 g dextrose and 15 g agar/L) or in yeast glucose medium (YGM; 10 g yeast extract/L, 30 g glucose/L) supplemented with appropriate antibiotics at 16°C with orbital shaking (100 rpm) for at least five days to obtain yeast-like conidiospores that were used for plant inoculation. To this end, conidiospores were collected by centrifuging the suspensions at 2,000 g for 5 min and adjustment to a final concentration of 10^7^ spores/mL with 0.5% Tween 20 for inoculation by brushing on adaxial and abaxial sides of primary leaves of 11-day-old wheat plants. The inoculated plants were covered in a plastic tent for two days to secure high humidity, after which the tent was opened in one-side.

Fungal biomass was measured with real-time PCR using a C1000 Touch™ Thermal Cycler (Bio-Rad, California, USA) with the *Z. tritici*-specific *β*-*tubulin* primers *Ztβtubulin*-F/R in combination with primers *TaCDC*-F/R that target the constitutively expressed cell division control gene of wheat (Table S1). Relative fungal biomass was calculated with the E^−ΔCt^ method and boxplots were made with RStudio using the package of ggplot2 (R Core Team, 2017; Wickham, 2016).

### Protein structure prediction and polymerization assay

Three-dimensional structures of Mgx1LysM was predicted with I-TASSER server (Roy *et al.*, 2010; Yang and Zhang, 2015). For chitin-induced polymerization assay of LysM effectors, concentrations of Mgx1LysM, Mg1LysM, RiSLM and Ecp6 were adjusted to 200 μM and 200 μL of each protein was incubated with 200 μL of 2 mM chitohexaose (Megazyme, Wicklow, Ireland), or 200 μL water as control, at room temperature overnight. The next day, 2 μL of 0.2% methylene blue (Sigma-Aldrich, Missouri, USA) was added and incubated for 30 min after which protein solutions were centrifuged at 20,000 g for 15 min. Photos were taken with a ChemiDoc MP system (Bio-Rad, California, USA) with custom setting for RFP.

## ACKNOWLEDGMENTS

H. Tian acknowledges receipt of a PhD fellowship from the China Scholarship Council (CSC). Work in the laboratory of B.P.H.J. Thomma is supported by the Research Council Earth and Life Sciences (ALW) of the Netherlands Organization of Scientific Research (NWO) and by the Deutsche Forschungsgemeinschaft (DFG, German Research Foundation) under Germany’s Excellence Strategy – EXC 2048/1 – Project ID: 390686111. J.J. Rudd and H. Chen were supported by the Biotechnology and Biological Sciences Research Council (BBSRC) of the UK Designing Future Wheat (DFW) Institute Strategic Programme (BB/P016855/1).

## AUTHORSHIP CONTRIBUTIONS

HT, LRM, JRM, BPHJT conceived the study; HT, CIM and LRM designed experiments; HT, CIM and GCMB performed experiments; HT analyzed data and wrote the manuscript; JJR and HC provided experimental materials, JRM and BPHJT supervised the project; all authors discussed the results and contributed to the final manuscript.

## DATA AVAILABILITY STATEMENT

Data sharing not applicable – no new data generated

