## Supplemental Table 1 for "Three LysM effectors of *Zymoseptoria tritici* collectively disarm chitin-triggered plant immunity"

**Table S1. Primers used in this study.**

| Primer name | Sequences |
| --- | --- |
| *Mgx1LysM*-userL-F | GGTCTTAAUAGAAAGGGCACTATTCAACAAGC |
| *Mgx1LysM*-userL-R | GGCATTAAUATACCATGCTGCCGAAGTTGAA |
| *Mgx1LysM*-userR-F | GGACTTAAUAAGGCGAAGCTCTTGAAAACTGG |
| *Mgx1LysM*-userR-R | GGGTTTAAUATCCAATCTTCACGTACCCGGTTTC |
| *Mgx1LysM*-F | CACGCCACGAAGACGATACCAT |
| *Mgx1LysM*-R | TTCAAGAGCTTCGCCTTTGGG |
| NAT-F | GTCACCAACGTCAACGCACCG |
| *Mgx1LysM*-out-R | ATCGGCGCTGTACAGAAATATGCATA |
| *Mgx1LysM*-cDNA-F | GGTGGTGAATTCCAGAACAACGCACAGTGTCG |
| *Mgx1LysM*-cDNA-R | GGTGGTGCGGCCGCTTATTATCAGCTGACATGTTTCTTCAAG |
| *TaCDC*-F | CAAATACGCCATCAGGGAGAACATC |
| *TaCDC*-R | CGCTGCCGAAACCACGAGAC |
| *Ztβtubulin*-F | AACGGTCGTTACCTCACCTG |
| *Ztβtubulin*-R | ACGTTGTTCGGAATCCACTC |
